# Signatures of criticality in efficient coding networks

**DOI:** 10.1101/2023.02.14.528465

**Authors:** Shervin Safavi, Matthew Chalk, Nikos Logothetis, Anna Levina

## Abstract

The critical brain hypothesis states that the brain can benefit from operating close to a second-order phase transition. While it has been shown that several computational aspects of sensory information processing (e. g., sensitivity to input) are optimal in this regime, it is still unclear whether these computational benefits of criticality can be leveraged by neural systems performing behaviorally relevant computations. To address this question, we investigate signatures of criticality in networks optimized to perform efficient encoding of stimuli. We consider a spike-coding network of leaky integrate-and-fire neurons with synaptic transmission delays and input noise. Previously, it was shown that the performance of such networks varies non-monotonically with the noise amplitude. Interestingly, we find that in the vicinity of the optimal noise level for efficient coding, the network dynamics exhibits signatures of criticality, namely, the distribution of avalanche sizes follows a power law. When the noise amplitude is too low or too high for efficient coding, the network appears either super-critical or sub-critical, respectively. Our work suggests that two influential, and previously disparate theories of neural processing optimization – efficient coding, and criticality – may be intimately related.

---

Attempts to understand information processing in the brain have led to the formulation of various optimality principles. Two major paths, among others, have been explored to uncover these principles. On one hand, a large body of studies starts from the theoretical and experimental finding that neural networks operate close to criticality (1, 2). Researchers have thus sought to investigate what, if any, could be the computational advantages of a network operating near a critical point (3). Meanwhile, another line of research presumes that neural networks have evolved to efficiently encode natural inputs (given constraints such as limited energy and noise). Here, the key question was investigating how neural networks could achieve such optimal encoding, and what are the resulting dynamics. In a nutshell: one line of research starts with an observation about neural dynamics (i.e., that they are near-critical) and seeks to find the coding advantages; the other starts with a coding objective (e.g., efficient coding) and seeks to understand the resulting dynamics. However, despite the prevalence of both approaches, connections between theories based on closeness to criticality and efficient coding hypothesis remain elusive.

To address this shortcoming, we introduce a complementary approach. Instead of tuning the network around the critical point and evaluating its statistical information processing performance, we optimize a network to perform a clearly defined computation and investigate if signatures of critical dynamics emerge in the optimized network. We focus on efficiently encoding the input, a well-established and functionally relevant computation, accompanied by a rich body of normative models (4) and neural dynamics (e. g., 5).

We analyzed a network of leaky integrate-and-fire (LIF) neurons that can be optimized (by adjusting the noise) to code a one-dimensional input (6). We evaluate the signatures of criticality, such as scale-freeness of the activity propagation cascades, termed neuronal avalanches, in networks with different noise amplitudes. Interestingly, we only observed scale-free neural avalanches in the vicinity of optimal noise for efficient coding. This result suggests that coding-based optimality co-occur with closeness to criticality.

## Results

We investigate a network of LIF neurons consisting of an excitatory and an inhibitory population. The network’s dynamics and connectivity is set up such that it can precisely encode a feed-forward input using a minimal number of spikes. In an idealized network with instantaneous synapses (7), recurrent inhibition removes redundancy between neurons. However, the introduction of realistic synaptic delays leads to network synchronization that impairs coding efficiency (for more details, see, e. g.,, 6). In the presence of synaptic delays, this network can nonetheless be optimized for efficient coding by adding noise (6, 8, 9) (or increasing the L1 norm which controls the spiking threshold, see 10). As it was shown in previous studies (see, e. g., 6, 15), the network’s performance depends non-monotonically on the noise amplitude, with the optimal performance achieved for an intermediate noise level.

To assess the signatures of criticality in the efficient coding network, we investigate the distribution of neural avalanches in networks with different levels of noise. To begin, we keep the network size fixed, at *N* = 100 (as originally used in, 6). A neuronal avalanche is defined as an uninterrupted cascade of spikes in the network (11). As suggested by (11), the period without spiking signifies the end of the previous avalanche if it is longer than the mean inter-spike interval (ISI) in the compound spike train (obtained by collapsing the spike trains of all neurons). Similar results were obtained using other thresholding choices (see supplementary methods), as in (11). In a system operating close to criticality, the avalanche size (number of spikes in the cascade) follows a power-law distribution. We demonstrate that the distribution of avalanche sizes systematically changes with the strength of added noise (Figure 1A). In networks with a small amount of noise (e. g., noise strength 0.5, thick blue line in Figure 1A, or Figure 1B left), large avalanches dominate the distribution of avalanche sizes (a bump in the tail of the distribution signifies a transient synchronization in the network). On the other hand, for a large amount of noise (e. g., noise strength 5.5, thick red line in Figure 1A, or Figure 1B right), the distribution is concentrated on the small avalanches (an exponential distribution). However, for intermediate levels of noise (e. g., noise strength 1.3, thick green line in Figure 1A, or Figure 1B middle), the avalanche-size distribution resembles a power-law (appears as a linear function in the log-log coordinates), which is a key signature of criticality in neural systems (see, e. g., 11).

**Fig. 1.**
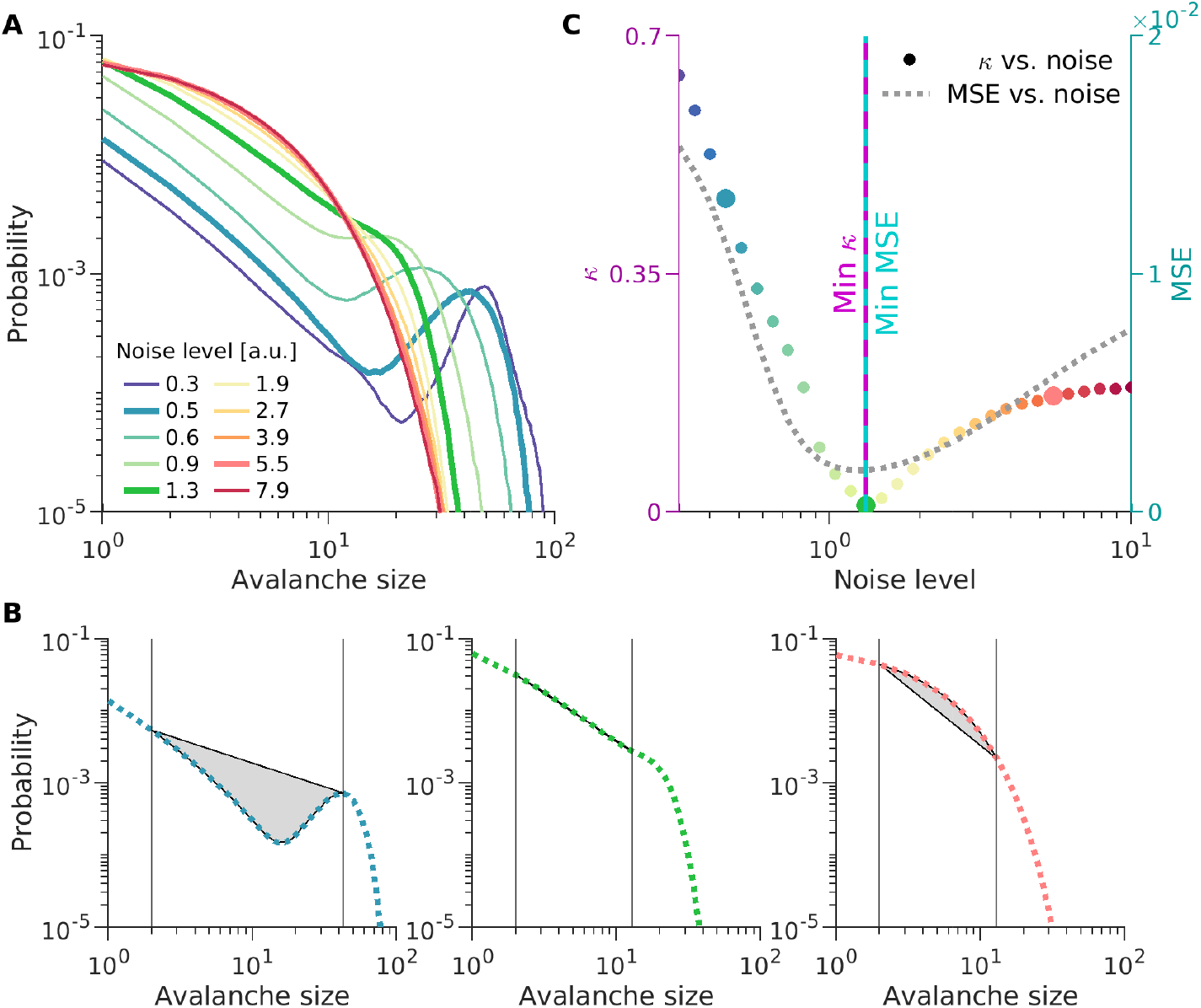
Co-occurrence of the criticality and optimal settings for efficient coding. **(A)** Avalanche-size distributions of efficient coding networks with different noise levels (indicated in the legend and with consistent color code across all panels). **(B)** Deviation from criticality measure *κ* for three noise levels. Left: small noise (0.5, network appears supercritical, many large avalanches); middle: medium noise (1.3, network close to criticality); right: strong noise (5.5, network exhibit subcritical behavior with predominantly small avalanches). (thicker lines in A): The area of gray-shaded regions between the actual avalanche size distribution and fitted power-law distribution defines the deviation measure *κ* (in the middle panel, the filled region is not visible, as the avalanche size distribution is very close to the ideal power-law). Vertical lines indicate the choices of left and right cut-offs (see main text for more details). Distributions for the chosen noise levels are highlighted in bold in panel A (matching colors). **(C)** Deviation from power-law *κ* as a function of noise level (left y-axis), color matching the panel A. Gray dotted line indicates mean-square-error (MSE) (right y-axis) as a function of noise. Vertical continuous line (purple) indicates the noise level corresponding to minimal *κ* (the most scale-free avalanche size distribution), Vertical broken line (cyan) indicates the noise level corresponds to minimum MSE (the best efficient coding performance). These two vertical lines overlap exactly, demonstrating the coincidence of noise levels for scale-free behavior and efficient coding.

To determine the most scale-free avalanche distribution (closest to a power-law distribution), we use a deviation measure *κ* that quantifies deviation from an ideal power-law distribution. Our *κ* measure closely follows the non-parametric measure introduced by Shew and colleagues (12) but does not assume a particular scaling exponent (see supplementary methods). *κ* is defined as the normalized area between the empirical and the ideal (fitted power-law for the portion of data between two cut-offs) distribution (Figure 1B). *κ* takes small (close to zero) values for a scale-free distribution (Figure 1B middle) and deviates from zero otherwise (Figure 1B left and right panels).

We measure how deviations from a power-law, *κ*, and the network’s reconstruction error depends on the noise strength. We confirm the previous observation that the performance of this network depends non-monotonically on the noise amplitude (gray dotted curve in Figure 1C), with the optimal performance achieved for an intermediate noise level (6, 15). Interestingly, the change in *κ* with the noise level demonstrates a similar non-monotonic behavior (colorful circles in Figure 1C). Remarkably, they both are minimized at the same noise level, resulting in a coincidence of the optimal point for coding and the most scale-free distribution (vertical purple and cyan line in Figure 1C). This observation offers additional support to the criticality hypothesis for the brain, namely that the various information processing measures are optimized close to the critical point (3, 13).

We next verify the stability of this result to changes in the network’s size by considering networks of various sizes in a range between *N* = 50 and *N* = 400 neurons. We find that all networks demonstrate similar non-monotonic behavior for the dependence of reconstruction error (Figure 2A) and the scale-freeness deviation measure *κ* (Figure 2B) on the strength of noise. This non-monotonic behavior of the reconstruction error is less pronounced for larger networks. This is expected, because the recurrent network used in our study is particularly suitable to code a single dimension of input by a small number of neurons(6, 7), i. e., hundreds, rather than thousands, of neurons per input dimension (see 10, as to how this problem could be alleviated for large networks by encoding higher dimensional inputs).

**Fig. 2.**
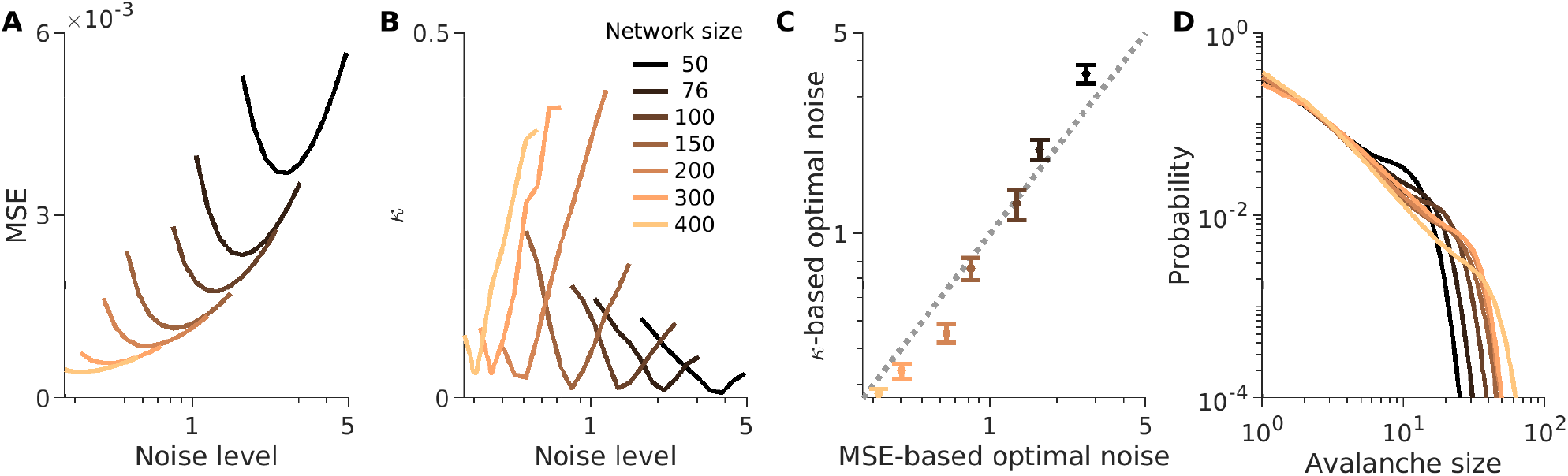
Co-occurrence of the criticality and efficient coding optimality across networks of different sizes. **(A)** Mean-square-error (MSE) of stimulus reconstruction for different injected noise amplitudes (similar to Figure 1C, gray line). Curves with different colors correspond to different network sizes (specified in the legend of panel B). **(B)** Deviation measure (*κ*) as function of noise (similar to Figure 1C, colorful dots). As in (A), different curves represent *κ*-noise relationship for networks of different sizes. **(C)** Y-coordinate of each point specifies the value of optimal noise chosen based on scale-freeness of the avalanche size distributions (minimum of *κ*), and the X-coordinate specifies the value of optimal noise chosen based on efficient coding criterion (minimum MSE). Error bars (mean ± standard deviation) indicate the variability across a wide range of choices of free parameters used for computing the deviation measure *κ* (see the supplementary text for more detail). **(D)** The cutoff of the power-law distribution for the most scale-free avalanche size distribution (resulting in smallest *κ* on panel B) for different network sizes shifts with the network size as expected from finite-size scaling ansatz for critical systems (colors are specified in the legend of panel B).

We observe the co-occurrence of efficient coding optimality with criticality optimality across all network sizes. The noise levels where coding error is minimal (x-coordinates in Figure 2C) and where *κ* is minimal (y-coordinates in Figure 2C) are highly correlated across different network sizes. Furthermore, this observation is robust to variations in the choice of the right cut-off needed for calculating *κ* (whiskers in Figure 2C indicate the standard deviation with respect to changing the ways *κ* is computed). Lastly, the location of the cut-off of the scale-free distribution shifts right (to the larger values) with the size of the network (Figure 2D), hinting at the correct finite-size scaling behavior (see, e. g., 11, 14).

## Discussion

In this study, we probe the connection between the optimality discussed in the context of criticality hypothesis of the brain, and the optimality discussed in theories of neural computations. To this end, we examine an efficient coding network (6, 7) for signatures of criticality. We find signatures of criticality (scale-free dynamics of the neural avalanches) emerging in a network that was *designed based on criticality-agnostic principles* merely by optimizing the coding performance. This suggests criticality and efficient coding are intimately related.

Our approach contrasts with previous work investigating the criticality hypothesis, which used models (e. g., a branching network, a recurrent neural network) that can attain various (critical/non-critical) states depending on a limited number of control parameters (e. g., branching ratio, connection strength) and then quantified how the computational primitives (3, 13), such as sensitivity to an input, depend on these control parameters. These state-generating models aim at reproducing realistic neuronal dynamics. They are typically driven by a slowly delivered noise and have no specific input and no readout strategy. Therefore, studies based on branching networks are largely agnostic to computational objectives central to theories of neural computation. Our approach thus paves the way for new questions about the relevance of criticality for precisely defined and task-relevant computations.

Future research should address why and how critical dynamics enables optimal efficient coding. For instance, it is not clear what is the exact role of neural avalanches, and why and how their scale-free distribution may be optimal for neural coding. Answering such questions requires going beyond our simulation-based approach, and, similar to (15). Mathematical analysis is needed to understand how the distribution of avalanches depends on different attributes of the network (noise, delay, connection weights, etc) and how those attributes affect the coding optimality. Such approaches will provide a mechanistic insight for our observations, and also, allow us to extend our framework to more sophisticated computations. We believe, our study opens up promising avenues for future investigations to establish the connection between other aspects of criticality (e. g., 16, 17) and theories of neural computations (e. g., 7).

## Materials and Methods

Further details are provided in the Supporting Information.

## ACKNOWLEDGMENTS

We thank Roxana Zeraati for sharing her codes for analysis of power-laws and her input on visualization; Victor Buendia for helpful discussions; Joachim Werner and Michael Schnabel for their IT support. This work was supported by the Max Planck Society, International Max Planck Research School for Cognive and Systems Neuroscience, and the Sofja Kovalevskaja Award from the Alexander von Humboldt Foundation to AL.

## Supplementary Information for

## Supporting Information Text

### Materials and methods

#### Efficient coding network

The neuronal network model used in this study was introduced and described extensively in the previous studies (1, 2), thus we restrict ourselves to a brief explanation of the key aspect of the model. Our network can be optimized to encode a sensory input efficiently (i. e., with a minimal number of spikes) and accurately (i. e., with minimal reconstruction error). Network optimization objective is incorporated in the loss function *E*(*t*),

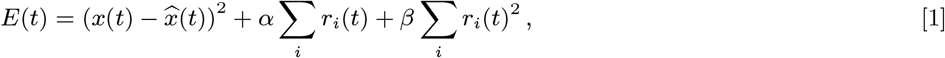

where *x*(*t*) is a given one-dimensional sensory input (similar to 2, 3), 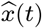 is the reconstructed sensory input, *r*_*i*_(*t*) is the firing rate of the neuron *i*, and *α* and *β* are the weights of the *L*1 and *L*2 penalties on the firing rate.

It is assumed that the input can be reconstructed by performing a linear readout of the spike trains, more precisely, by a weighted leaky integration of output spike trains,

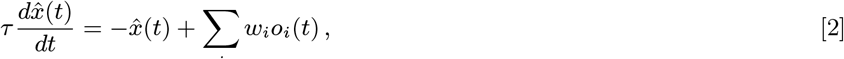

where *o*_*i*_ indicates the output spike trains for the neuron *i*,

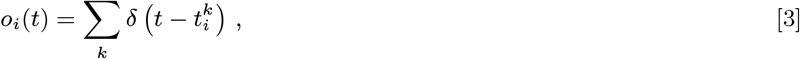

and *τ* is the read-out time constant^∗^, and *w*_*i*_ is a constant read-out weight associated to the neuron *i*.

Given an idealized network with instantaneous synapses, the optimal network could be derived from first principles. Boerlin *et al*. (1) demonstrated that the dynamics of each leaky-integrate and fire (LIF) neuron can be expressed by conventional differential equation governing the dynamics of the membrane potentials,

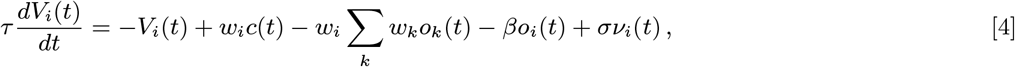

where *V*_*i*_ is the membrane potential of the neuron *i, w*_*i*_ is the constant readout which was introduced in Equation 2, *c*(*t*) is the input to the network, *o*_*i*_(*t*) is the spike train of neuron *i, β* is the regularizer that was introduced in Equation 1, and *ν*(*t*) is a white noise with unit variance that was manually added in the original derivation of (1) for biological realism. Notably, in this network we have two types of input, a feed-forward input, *w*_*i*_*c*(*t*) and a recurrent input −*w*_*i*_ *Σ*_*k*_ *w*_*k*_*o*_*k*_(*t*). The recurrent input is the result of a fully connected network. In this network, neurons that receive a common input, decorrelate their activity to avoid communicating redundant information via instantaneous recurrent inhibition.

Chalk *et al*. (2) introduced a more biologically plausible variants of (1)’s network by incorporating synaptic delays and introducing a balance network of inhibitory and excitatory population of neurons. They incorporated realistic synaptic delays by assuming that each spike generates a continuous current input to other neurons, with a dynamic that is described by the conventional alpha function,

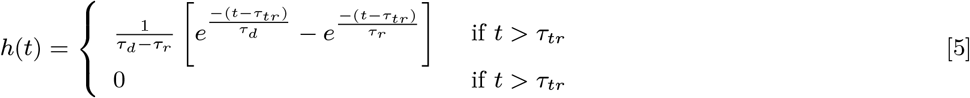

where *τ*_*r*_ and *τ*_*d*_ are respectively synaptic rise and decay times. Adding realistic synaptic delays, led to network synchronization, which impairs coding efficiency. Chalk *et al*. (2) demonstrated that, in the presence of synaptic delays, this network of LIF neurons can nonetheless be optimized for efficient coding by adding noise to the network. In this study, we implement the additional noise, as white noise added to the membrane potentials. However, (2) also demonstrated similar dependency of network’s performance to noise by using other ways of incorporating noise, for instance, by inducing unreliability in spike elicitation (also see, 4–6, for other approaches).

The original network introduced by (1) was a pure inhibitory network. (2) introduced a variant of this network that respects the Dale’s law. In their network, they introduce a population of inhibitory neurons that tracks the estimate encoded by the excitatory neurons, and provides recurrent feedback to the excitatory population (for further detail see, 1, 2)

#### Avalanche detection

To investigate the scale-free characteristic of the spiking activity (as a potential signature of networks operating close to criticality), similar to previous studies (7), we probe the distribution of neural avalanches. A neuronal avalanche is defined as an uninterrupted cascade of spikes in the network (7). In a system operating close to criticality, the distribution of avalanche sizes (number of spikes in a cascade) and avalanche life-time follows a power-law (in this study we have only investigated the distribution of avalanche sizes).

For detecting the avalanches, we followed the procedure used in previous studies (e. g., 7). The period of no spiking activity signifies the end of the previous avalanche if it is larger than a threshold ∆. We mainly used as a threshold the mean inter-spike interval (∆ = ⟨*ISI*⟩) in the compound spike train (obtained by collapsing the spike trains of all neurons onto a single time-line). Thus, when the compound spike train is interrupted for an interval larger than ∆, we consider that the current avalanche is over, and a next spike will be an onset of a new avalanche. The size of the avalanche is the number of spikes between these two silent time-points. A slightly different procedure has also been used for avalanche detection. In the alternative approach, for computing the ∆, one counts the synchronous spike only once, i. e., excluding zero *ISI*s. Notably, similar results were obtained using the alternative method.

This choice of threshold ∆ potentially can be made separately for individual network with different noise levels. However, to avoid introducing an additional variability across different levels of noise, we fixed the threshold for all the noise levels. For a given network size, we took the threshold from the network with the noise level corresponding to minimal mean-square-error (MSE), and use that for all noise levels. We also checked that taking slightly different thresholds would not change the results of our study.

#### Closeness to criticality assessment

We consider the scale-free distribution of neural avalanche as signature of criticality in the network (8, 9). Thus, to determine the most scale-free avalanche distribution, we introduce a deviation measure *κ*, which quantifies deviation from an ideal power-law distribution. Our *κ* measure closely follows the non-parametric measure introduced by Shew *et al*. (10), but does not assume a particular scaling exponent, which might be important, because we do not know *a priori* what is the relevant universality class for neuronal avalanches (see, e. g., 11). We define *κ* as the area between the empirical and the ideal (fitted power-law) distribution, normalized by the number of data-points (in the empirical distribution) between left and right cut-offs. Larger values of *κ* correspond to larger deviations from the power-law distribution. Power-laws were fitted between two cut-offs. The left cut-off was always chosen to be 2 (i. e., avalanches with at least two spikes). The right cut-off is typically chosen subjectively based on the problem at hand (12), here we swept over a wide range of choices to be as objective as possible. We choose it between two possible options: it was either a certain percentile of number of avalanches (within the range of 50-95%), or a fraction of the network size (within the range of 10-25%). Between the mentioned choices above, the one led to inclusion of more data, i. e., larger proportion of avalanches was selected The ideal power-law distribution were also determined based on a linear fit between final choices of left and right cut-offs. Notably, the results were robust to variations (in the ranges noted above) in the choice of cut-offs (see Figure 2C of the main text).

## Footnotes

∗ In the efficient coding network used in this study (as in 2), for simplicity, the read-out time constant of the input (i. e., time-scale of *x*(*t*)) is the same as the time-constant of the membrane potential of the neurons. Nevertheless, in (1) they are not necessarily the same for more general computations.

